# Heparan sulfates regulate axonal excitability and context generalization through Ca^2+^/calmodulin-dependent protein kinase II

**DOI:** 10.1101/2023.01.26.524131

**Authors:** Inseon Song, Tatiana Kuznetsova, David Baidoe-Ansah, Hadi Mirzapourdelavar, Oleg Senkov, Hussam Hayani, Andrey Mironov, Rahul Kaushik, Michael Druzin, Staffan Johansson, Alexander Dityatev

## Abstract

Our previous studies demonstrated that enzymatic removal of highly sulfated heparan sulfates with heparinase 1 impaired axonal excitability and reduced expression of ankyrin G at the axon initial segments in the CA1 region of the hippocampus ex vivo, impaired context discrimination in vivo, and elevated Ca2+/calmodulin-dependent protein kinase II (CaMKII) activity in vitro. Here, we show that in vivo delivery of heparinase 1 in the CA1 region of the hippocampus elevated autophosphorylation of CaMKII 24 hours after injection in mice. Patch-clamp recording in CA1 neurons revealed no significant heparinase effects on the amplitude or frequency of miniature excitatory and inhibitory postsynaptic currents, while the threshold for action potential generation was increased and fewer spikes were generated in response to current injection. Delivery of heparinase on the next day after contextual fear conditioning induced context overgeneralization 24 hours after injection. Co-administration of heparinase with the CaMKII inhibitor (autocamtide-2-related inhibitory peptide) rescued neuronal excitability and expression of ankyrin G at the axon initial segment. It also restored context discrimination, suggesting a key role of CaMKII in neuronal signaling downstream of heparan sulfate proteoglycans and highlighting a link between impaired CA1 pyramidal cell excitability and context generalization during recall of contextual memories.

## 1. Introduction

Heparan sulfate proteoglycans (HSPGs) are harboring long chains of variously sulfated polysaccharide residues. There are membrane-bound HSPGs, such as syndecans and glypicans, and secreted HSPGs, including agrin, perlecan and collagen type XVIII. Accumulating studies demonstrate that HSPGs have an important role in the nervous system during development and adulthood.

In the mouse brain, syndecan-1 and glypican-4 are highly expressed in the neural tube, where the precursor cells are proliferating [1]. These HSPGs are important for the proliferation of neural precursor cells and play a role as synaptic organizing molecules during synaptogenesis. Their HS chains are essential for this role. Glypican 4 is bound to the presynaptic membrane via a GPI anchor and interacts with the postsynaptic protein, LRRTM4 (leucine-rich repeat transmembrane neuronal proteins), forming a trans-synaptic complex. This complex recruits other synaptic molecules to the synaptic cleft, contributing to the maturation of excitatory synapses. Mice deficient in glypican 4 exhibit decreased synapse number along with decreased postsynaptic GluA1 and increased retention of presynaptic neuronal pentaxin 1 [2].

Syndecans are differentially expressed in various neural cell types and exhibit differential subcellular localization in neuron [3]. Contrary to glypicans lacking a cytoplasmic domain, transmembrane syndecans interact with specific cytoplasmic binding partners such as CASK, syntenin, synectin, and synbindin [4-7]. Syndecan 2 is highly expressed in synapses and influences activities of postsynaptic scaffolding protein, thereby contributing to filopodia and dendritic spine formation [8]. Overexpression of full-length syndecan 2 in cultured immature hippocampal neurons accelerates dendritic spine formation, while a syndecan 2 deletion mutant that lacks the ability for binding to synthenin and CASK does not support spine maturation [4,9]. Association of cortactin and fyn to syndecan is increased rapidly after induction of long-term potentiation (LTP), while inclusion of soluble syndecan 3 into the rat hippocampal slices inhibits high-frequency stimulation-induced LTP [10]. Furthermore, syndecan 3 knockout mice exhibit strongly enhanced LTP and impaired hippocampal-dependent memory [11]. Secreted HSPG agrin is also involved in filopodia and dendritic spine formation. While downregulation of agrin in the cultured neurons *in vitro* and *in vivo* reduces the number of dendritic filopodia, overexpression of agrin in rodent hippocampal neurons stimulates filopodia formation *in vitro* [12].

Accumulating structural, pharmacological and genetic studies suggest a key role of HS chains carried by HSPG in mediating their activities. Interestingly, HSs bind to receptor protein tyrosine phosphatase sigma (RPTPσ) at the same site as chondroitin sulfates. Crystallographic analyses of this site reveal conformational plasticity that can accommodate diverse glycosaminoglycans with comparable affinities. HSs induced RPTPσ ectodomain oligomerization, stimulating neurite outgrowth. The oligomerization was inhibited by chondroitin sulfates, resulting in impaired neurite outgrowth [13]. In acute hippocampal slices, treatment with a mixture of heparinases 1 and 3, which removes highly and low sulfated HSs, respectively, impaired long-term potentiation (LTP) of synaptic transmission [10,14]. This treatment also prevented the increase in the number of spines after induction of NMDA receptor-dependent LTP [14]. Conditional ablation of Ext1, a gene involved in HS synthesis, in a subpopulation of pyramidal neurons leads to an autistic phenotype [15], providing genetic evidence for the importance of HSs to shape brain function on many levels from cellular properties to complex behaviors. More targeted ablation of HSs on neurexin-1 also revealed structural and functional deficits at central synapses. HS directly binds postsynaptic partners neuroligins and LRRTMs [16].

Considering the high heterogeneity of HSs, we focused on a highly sulfated subset of HSs (HSHSs), which could be digested by heparinase 1. Such treatment of cultured hippocampal neurons resulted in a reduction in the mean firing rate of neurons [17,18], despite the upregulation of GluA1 protein expression [17]. Acute treatment of hippocampal slices with heparinase 1 reduced CA1 pyramidal cellular excitability and impaired hippocampal LTP [19]. Altered expression of ankyrin G (AnkG) as one of the major organizing proteins at the axon initial segment (AIS) in heparinase 1-treated hippocampal slices led us to the hypothesis that HSHSs are involved in modulation of neuronal activity through the changes in the AIS composition and function. Injection of heparinase 1 before fear conditioning impaired context discrimination [19], validating the importance of HSHSs at the systemic level.

Based on previous in vitro findings of increased autophosphorylation levels of CaMKII α and β isoforms after heparinase 1 treatment [17], we hypothesized that CaMKII is the key molecule involved in the modulation of axonal excitability due to a loss of HSHSs and provided *in vitro* and *in vivo* evidence verifying this hypothesis biochemically, immunocytochemically and electrophysiologically. Our studies show that an increased level of autophosphorylated CaMKII in heparinase-treated neurons is responsible for reduced neuronal excitability, altered expression of AnkG in the AIS of CA1 pyramidal neurons, and impaired contextual discrimination.

## 2. Materials and Methods

### 2.1. Immunoblot analysis

To access the level of endogenous CaMKII isoform expression and the level of their phosphorylation, murine hippocampal slices (treated with intact or heat-inactivated heparinase 1 in the same way as for electrophysiological recordings) were snap-frozen in isopropanol pre-cooled on dry ice. Later samples were homogenized in RIPA buffer (ThermoFisher Scientific) containing a protease inhibitor cocktail (Sigma-Aldrich P1860), serine/threonine phosphatases inhibitor (Sigma-Aldrich P0044) and tyrosine phosphatases inhibitor (Sigma-Aldrich P5726) using glass tissue homogenizer. Non-soluble proteins were separated by centrifugation at 20,000 g for 15 min at 4 °C. The protein concentration of individual samples was measured using DC Protein Assay (Bio-Rad). 10-30 μg of extracts were resuspended in reducing (5.0 % 2-mercaptoethanol) sample buffer (Bio-Rad) and boiled at 100°C for 5 min, separated by SDS-PAGE on 10% acrylamide gels and transferred to the PVDF membranes. Membranes were blocked for 1h at room temperature with 5% Blotting-Grade Blocker (Bio-Rad, 1706404) in TBS-T buffer, probed with appropriate primary antibody at 4°C overnight and subsequently for 1 h at room temperature with HRP-conjugated secondary antibodies.

To estimate the total expression of α and β forms of CaMKII mouse anti-CaMKII (G1, sc-5306; 1:200 -1:1000) from Santa-Cruz was used. To access activation of CaMKII, rabbit anti-phospho Thr 286/287 CaMKII (p1005-286; 1:1000) from PhosphoSolutions was applied. To evaluate the GAPDH level, mouse anti-GAPDH (MAB374; 1:15.000-20.000) from Millipore was used. HRP-conjugated secondary antibodies were donkey anti-rabbit (NA934V) from GE Healthcare, or goat anti-mouse (115-035-146) from Jackson ImmunoResearch. Acquisition of chemiluminescent signal and densitometric analysis were performed by Odyssey Fc imaging system (LI-COR) and Image Studio software respectively. The total levels of α or β forms of CaMKII and phospho Thr 286/287 level were standardized to the level of loading control (GAPDH) in each sample. Standardized values were further normalized to the randomly chosen control sample (loaded in each gel). To evaluate CaMKII phosphorylation on Thr 286/287, the ratios between phospho-Thr 286/287 signal to the total amount of CaMKII protein were calculated.

For statistical evaluation and the graphical representation of the data, the Origin software was used. The average ± SEM (standard error of mean) was calculated for control and experimental (heparinase-treated) groups, normalized to randomly chosen control samples. Statistical evaluation was carried out by Mann–Whitney–Wilcoxon test.

### 2.2 Slice preparation and in vitro electrophysiology

Acute hippocampal slices were prepared as described elsewhere [19] from 4- to 5- week-old male C57Bl/6 mice 1 day after injection of Heparinase 1, or ctrl (heat-inactivated Heparinase 1), or Heparinase 1 + autocamtide-2-related inhibitory peptide (AIP) into hippocampal CA1 area as described below (*Intrahippocampal injection)*. Transverse 350 μmthick hippocampal slices were obtained in ice-cold slice solution containing (in mM) 240 sucrose, 2 KCl, 2 MgSO_4_, 1.25 NaH_2_PO_4_, 26 NaHCO_3_, 1 CaCl_2_, 1. MgCl_2_, and 10 D-glucose. After slice recovery at room temperature, the slices were transferred to a submerged recording chamber and were perfused with ACSF (2-3 ml/min) containing (in mM) 124 NaCl, 2.5 KCl, 1.3 MgSO_4_, 1 NaH_2_PO_4_, 26.2 NaHCO_3_, 2.5 CaCl_2_, 1.6 MgCl_2_, and 11 D-glucose the solution was saturated with 95% O_2_/5% CO_2_ (Osmolarity, 290 ± 5 mOsm). Wholecell patch-clamp recordings were obtained from visually identified CA1 pyramidal neurons with glass electrode (4-5 MΩ, Hilgenberg, Germany) containing (in mM) 120 K-gluconate, 10 KCl, 3 MgCl_2_, 0.5 EGTA, 40 HEPES, 2 MgATP, 0.3 NaGTP (pH 7.2 with KOH, 295 mOsm) for measuring neuronal excitability. In the current clamp configuration, cells were held at -70 mV and injected from -75 mV to +400 pA with 25 pA increments. For measuring mEPSCs, 5 mM QX314 was added into the intracellular pipette solution while GABA_A_ receptor antagonist picrotoxin (PTX, 50 μM, Tocris), GABA_B_ receptor antagonist CGP55845 (3 μM, Tocris), Na^+^ channel blocker tetrodotoxin (TTX, 1 μM, Tocris) were added to ACSF.

Miniature IPSCs were recorded with a glass electrode containing (in mM) 120 CsCl, 8 NaCl, 0.2 MgCl_2_, 10 HEPES, 2 EGTA, 0.3 Na_3_GTP, 2 MgATP, pH 7.2 with CsOH, 290 mOsm). NBQX (25 μM,Tocris), D-APV (50 μM,Tocris), TTX (1 μM, Tocris) were added to ACSF to isolate action potential-independent mIPSCs. *In vitro* electrophysiological data were acquired using an EPC 10 amplifier (HEKA Elektronik, Germany) at a sampling rate of 10 kHz, low pass filtered at 2-3 kHz. The obtained data were offline analyzed using PatchMaster software (HEKA Electronik, Germany), pClampfit 10 (Molecular Devices, U.S.A.), or MiniAnalysis (6.0.3 Synaptosoft, U.S.A.). The data were presented and analyzed using Sigmaplot 12 (Systat Software Inc, U.S.A.) and Prism 7 (Graphpad software, U.S.A.).

### 2.3. Immunocytochemistry in hippocampal cultured neurons

Hippocampal neurons from embryonic C57BL6/J mice (E18) were extracted and cultured as described earlier [19]. The neuronal cells were plated on polyethyleneimine-coated (Sigma-Aldrich; 408727-100ml) 18 mm coverslips in 12 well plates with a cell density of 150,000 per well. Neurons were maintained in 1 ml of Neurobasal media (NB+ media) (Invitrogen) containing 2% B27 and 1% L-glutamine and 1% Pen-Strep (Life Technologies). Cultured neurons were fed with 250 μl NB+ media on days *in vitro* (DIV) 14 and 17. On DIV 21-23, cultured hippocampal neurons were incubated with Heparinase-1 (0.5 U/ml, Sigma-Aldrich, H2519), Ctrl (heat-inactivated Heparinase-1), or Heparinase-1 + AIP (100 nM) as described [19] for 2.5 h at 37 °C. After the treatment, hippocampal neurons were washed with phosphate-buffered saline (PBS) and fixed with 4% paraformaldehyde (PFA) for 10 minutes, then permeabilized with 0.1% Triton-X-100 in PBS for 10 min, washed 3 times, and blocked (0.1% Glycine + 0.1% Tween-20 + 10% Normal Goat Serum in PBS) for 60 min at room temperature. Then, the cells were stained with antibodies against AnkG (mouse monoclonal, 1:1000; Millipore, MABN466), pCaMKII (rabbit polyclonal, 1:1000; Phospho Solution, P-1005-286), MAP2 (chicken polyclonal, 1:500; Abcam, ab5543) and DAPI (Life Technologies, S36939), and finally mounted (Fluoromount; Sigma Aldrich, F4680-25ML) for imaging. Mounted coverslips were imaged using Zeiss LSM 700 confocal microscope with a 63x/1.4 NA oil immersion objective. Image analysis was done as previously described [19]. Using the MAP2 and AnkG signals, the AISs were analyzed from the soma edge over a 40 μm long distance with a line profile (width=3.0) using Fiji [19].

### 2.4. In vivo intrahippocampal injection and fear conditioning

*Animals*. Adult (2- to 4-month-old) male C57Bl/6j mice (Charles River) were used. At least 1 week before starting the experiments, mice were transferred to a small vivarium, where they were housed individually with food and water *ad libitum* on a reversed 12:12 light/dark cycle (light on at 9:00 p.m.). All behavioral experiments were performed in the afternoons during the dark phase of the cycle when mice are active, under constant temperature (22 ± 1°C) and humidity (55 ± 5%). All treatments and behavioral procedures were conducted in accordance with ethical animal research standards defined by German law and approved by the Ethical Committee on Animal Health and Care of the State of Saxony-Anhalt, Germany, the license numbers 42502-2-1159 and -1322 DZNE.

*Intrahippocampal injection*. Injection guide cannulas and electrodes were implanted as previously described [19] but electrophysiological analysis is not included in the present study due to an insufficient quality of recordings. Coordinates for bilateral cannulas were: AP = 2.0mm and L = ±2.2mm from Bregma and midline, respectively. For intrahippocampal injection, we used a digitally controlled infusion system (UltraMicroPump, UMP3, and Micro4 Controller, WPI, USA) fed with a 10 μl Hamilton syringe and NanoFil (35 GA) bevelled needle as previously described [19]. The mouse was first anesthetized with 1-3% isoflurane and put into the stereotaxic frame. Heparinase 1 from *Flavobacterium heparinum* (0.05 U/μl/site, Sigma-Aldrich, H2519), or ctrl (heat-inactivated Heparinase 1 at 100 ºC for 30 min), or Heparinase 1 + autocamtide-2-related inhibitory peptide (AIP, 0.17 μg/μl/site, Sigma-Aldrich, SCP0001) was injected into hippocampal CA1 area at a rate of 3 nl/s. After waiting for another 5 min, the injection needle was removed.

*Fear conditioning*. In this study, we used the previously described classical Pavlovian contextual fear conditioning (CFC) paradigm in mice with slight modification [19]. Here, on day 0 (d0) mice were initially placed in a 20×20×30 cm chamber with a neutral context (CC-); grey walls, and grey plastic floor, for 5 min. Next, mice were exposed to the conditioned context (CC+) which includes patterned walls and a metal grid floor, for 5 min after an interval of 1 hour. During the CC+ phase, mice ‘s feet were shocked 3x with mild intensity (0.5 mA, 1 s) with a 1 min inter-shock interval. Using a computerized fear conditioning system (Ugo Basile, Italy), the first memory retrieval session was carried out for 5 min for each mouse on day 1 (d1) with a 1-hour interval following the sequence CCand CC+. On day 2 (d2), mice were injected with vehicle, Heparinase and Heparinase + AIP into the hippocampus. Then, on day 3 (d3) the second memory retrieval test (test 2) was performed using a similar paradigm as on d1. Additionally, a blinded trained observer used video recordings of each session for offline fear-conditioned behavioral analysis with the help of the behavioral video acquisition and analysis software (AnyMaze, USA). Finally, the overall context memory and discrimination performance for each mouse was estimated. *Statistics*

Numerical data are reported as mean ± SEM with n being the number of samples. Student ‘s t-test and multi-way ANOVA with suitable posthoc tests were used as indicated and performed in SigmaPlot or Prism. For non-Gaussian distributions, we used Mann– Whitney–Wilcoxon test. Significance levels (p-values) are indicated in figures by asterisks.

## 3. Results

### 3.1 Heparinase treatment elevates CaMKIIβ autophosphorylation in the mouse hippocampus

We have previously observed an increase in the GluA1 expression and CaMKII activity in cultured mouse hippocampal neurons after heparinase 1 treatment [17]. To investigate whether heparinase treatment also changes hippocampal CaMKII activity *in vivo*, Ctrl (heat-inactivated heparinase) or active heparinase 1 was injected into the dorsal hippocampus of 6-week-old mice. To investigate the level of endogenous CaMKII isoform protein expression and their activity, 24 hours after injection of heparinase hippocampal slices were acutely prepared and used for immunoblotting. CaMKIIα and CaMKIIβ are major isoforms in the hippocampus and these molecules are activated during memory formation. Activation of CaMKIIα and CaMKIIβ was assessed by the analysis of phosphorylation at Thr286 and Thr287, respectively [20]. Consistent with the previous observation in the cultured hippocampal neurons [17], the activity of CaMKIIβ was strongly affected after heparinase injection *in vivo*. The ratio between phosphorylated and total CaMKIIβ was increased after heparinase treatment, while the effect on CaMKIIα was less prominent (Figure 1).

**Figure 1.**
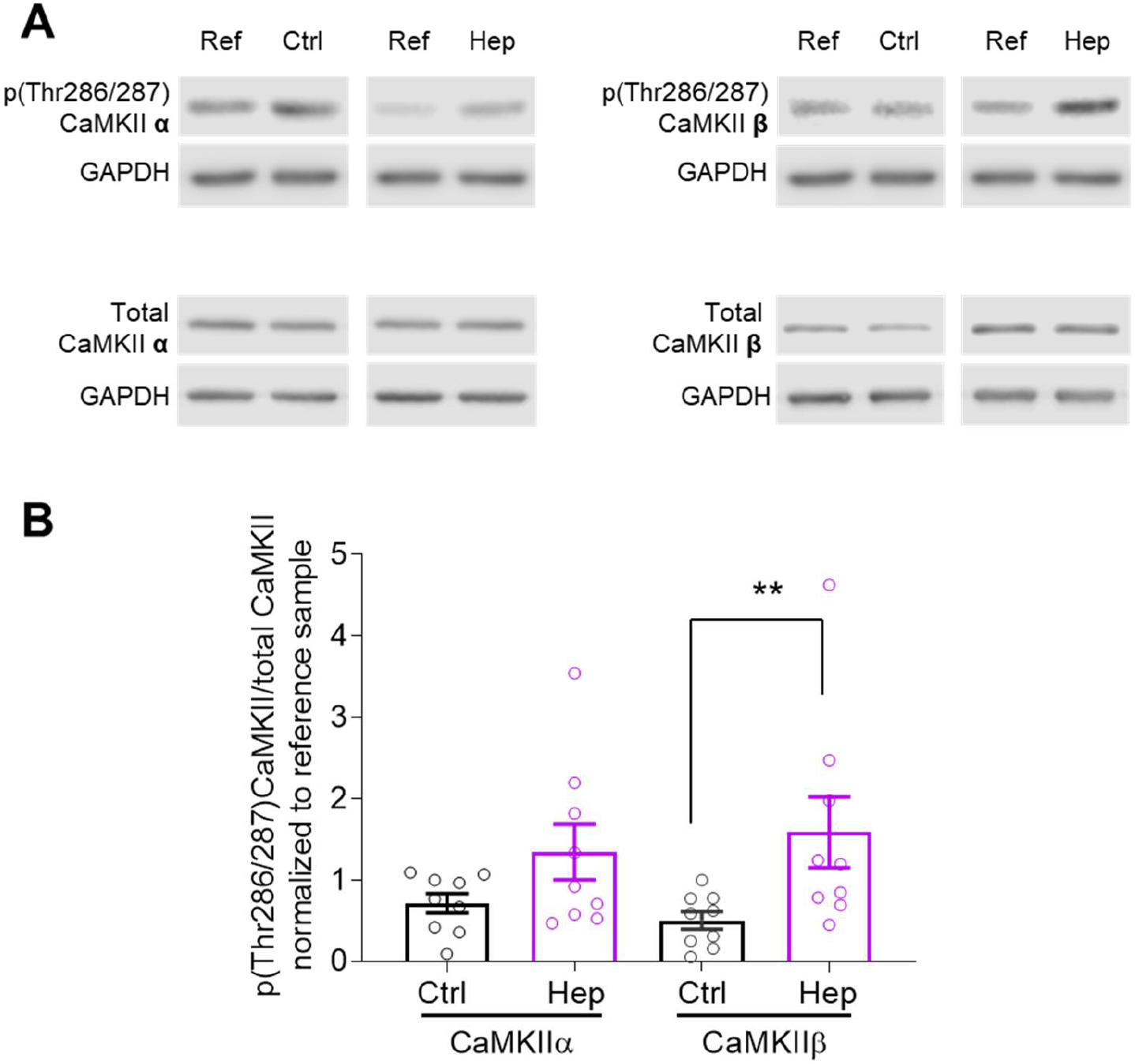
Heparinase treatment increased CaMKII activity in vivo. (**A**) Heparinase-injected mouse hippocampi were lysed and extracts were used for immunoblotting. Membranes were incubated with anti-phospho-Thr286/287 CaMKII antibody to measure autophosphorylated CaMKII α and β levels relative to the total protein expression levels. (**B**) Summarized graph showing the statistical evaluation of western blotting experiments as in (A). Data are presented as means ± SEMs. Note that there is a significant increase in CaMKIIβ autophosphorylation one day after heparinase injection (**p < 0.01, Mann–Whitney–Wilcoxon test; ctrl, n=9; Hep, n=9).

### 3.2. Enzymatic digestion of HSHSs does not change synaptic transmission to CA1 pyramidal neurons

Having found that heparinase treatment *in vitro* can up-scale mEPSCs, we measured glutamatergic transmission (mEPSC) and GABAergic transmission (mIPSC) to CA1 pyramidal neurons 1 day after heparinase injection *in vivo*. Unexpectedly, we observed changes neither in the mEPSCs amplitude nor the frequency (Figure 2A, C). Also, temporal parameters such as the rise and decay times were not affected (Figure 2A, C). The properties of mIPSCs were also unchanged by heparinase treatment (Figure 2B, D).

**Figure 2.**
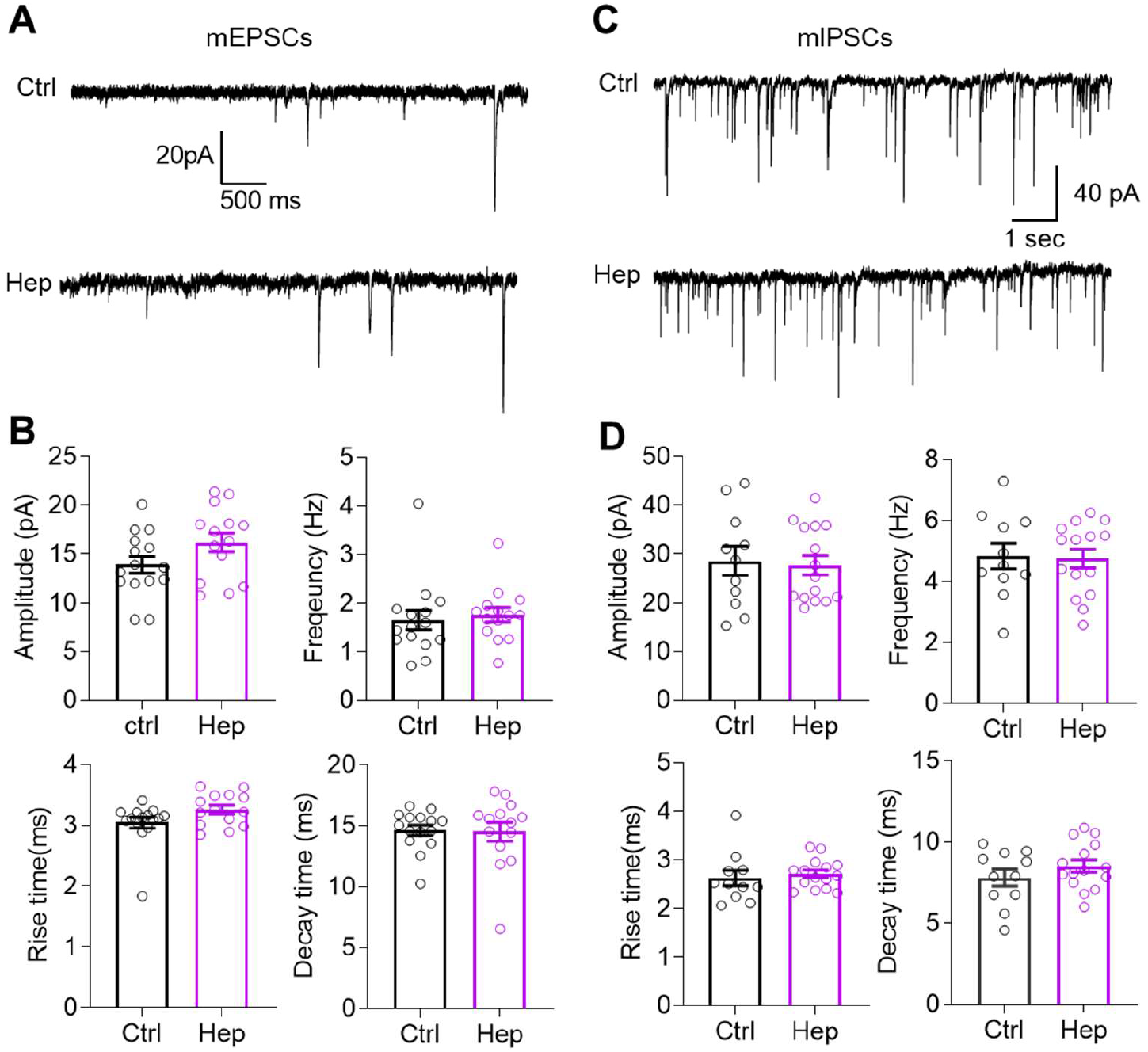
Intact excitatory and inhibitory synaptic transmission onto CA1 pyramidal cells one day after heparinase injection into the mouse hippocampal CA1 in vivo. (**A**) Representative traces of mEPSCs from CA1 pyramidal cells in heat-inactivated (Ctrl) and active heparinase 1injected mice in the presence of PTX, CGP55845, and TTX. (**B**) Summarized bar graphs show that amplitude, frequency, rise time, and decay time of mEPSCs remained intact after heparinase injection (p>0.05, two-tailed t-test; Ctrl, n=15; Hep, n=14). Data are presented as means ± SEMs. (**C**) Representative traces of mIPSCs recorded from CA1 pyramidal cells in heat-inactivated (Ctrl) and active heparinase 1-injected mice in the presence of NBQX (25 μM), D-AP5 (50 μM), CGP55845 (3 μM), and TTX (1 μM). (**D**) Summarized bar graphs show that amplitude, frequency, rise time, and decay time of mIPSC remained intact after heparinase injection into the hippocampal CA1 area (p >0.05, two-tailed t-test; Ctrl, n=15; Hep, n=14). Data are presented as means ± SEMs.

### 3.3. Impaired neuronal excitability after in vivo injection of heparinase is rescued by CaMKII inhibitor AIP

Next, we investigated the excitability of CA1 pyramidal neurons. We previously reported that acute heparinase treatment of hippocampal slices reduced AP probability during TBS and hence decreases Ca^2+^ influx to dendritic spines during induction of LTP [19]. Based on that study, we expected that one-day heparinase treatment may also result in reduced neuronal excitability in the CA1 pyramidal cells. Therefore, we performed patchclamp recordings in the current-clamp configuration and measured the number of action potentials (APs) as a function of injected currents (Figure 3A), the threshold of action potential generation (Figure 3B), and other parameters characterizing the magnitude and shape of APs (Figure 3C-F). To verify the role of CaMKII in the effects of heparinase, we employed AIP as a selective and potent inhibitor of CaMKII, which has been used in slices and *in vivo* [21-23]. We co-injected AIP with heparinase one day before recordings.

**Figure 3.**
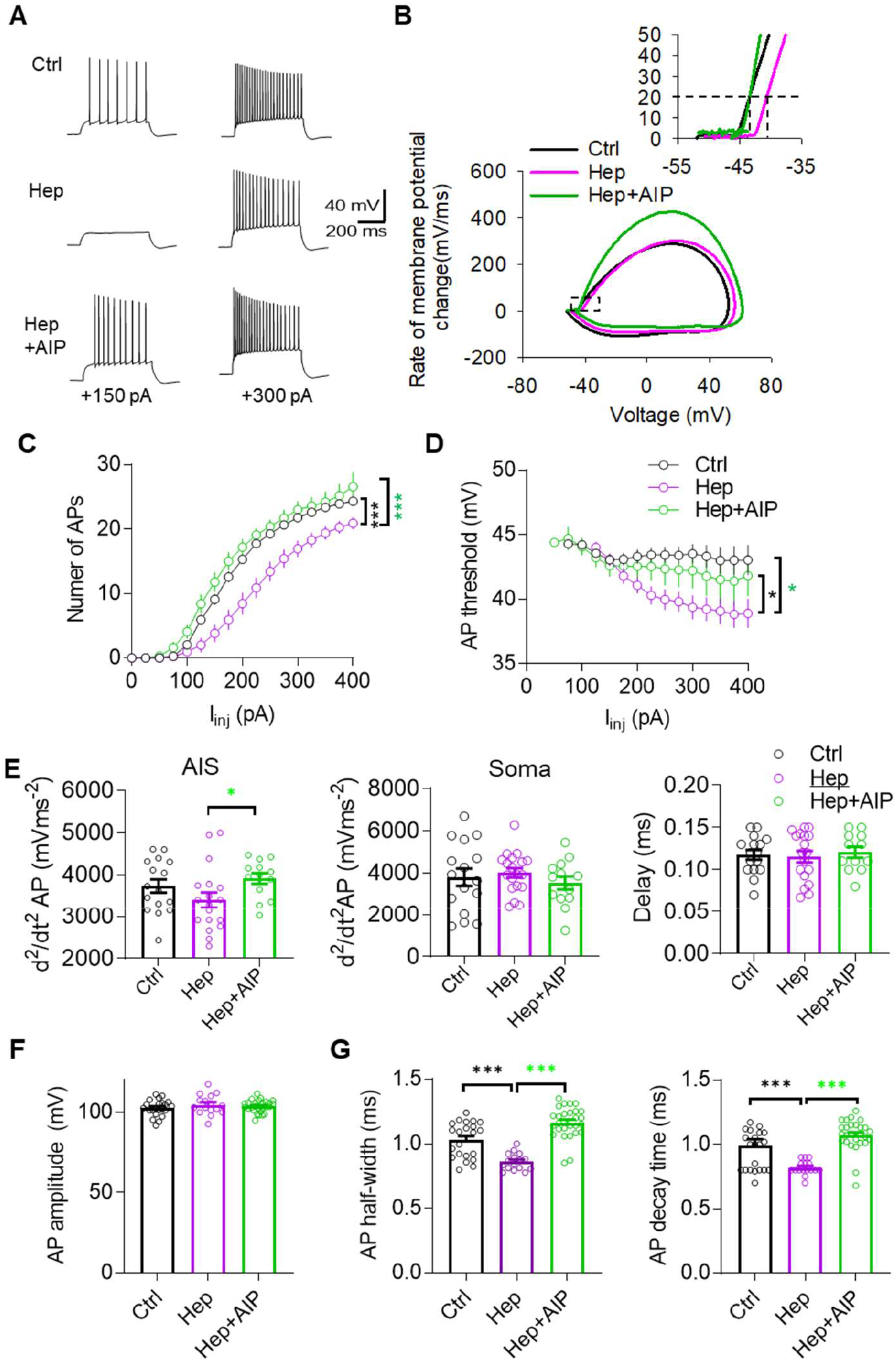
CaMKII inhibitor (AIP) could rescue reduced neuronal excitability of CA1 pyramidal neurons after heparinase injection in vivo. (**A**) Sample traces of action potentials (APs) recorded in the CA1 PC in response to depolarizing current injections. (**B**) Phase plot (dV/dt versus V) of the first action potential in response to 300 pA current injection. Inset (left) shows the initial phase of the action potential from the phase plot (right). (**C & D**) Summary graphs showing the number of action potentials generated in PC in response to different depolarizing current injections and the AP threshold at 20 mV/ms in response to different current injections. * p< 0.05, ***p<0.001, Holm-Sidak test after two-way RM ANOVA. (**E**) Analysis of the amplitude and interval (Delay) between two peaks of the second action potential derivative (d/dt2 AP), which correspond to the activation of sodium channels in the AIS and soma, revealed a facilitated AP generation at the AIS after heparinase treatment with co-administration of AIP. * p< 0.05, Holm-Sidak test after one-way ANOVA. (**F & G**) Summary graphs comparing Ctrl (heat-inactivated heparinase), heparinase, and heparinase+AIP-injected neurons in terms of the amplitude (E), HW (half-width), rise time, and decay time of the action potential (ctrl, n=22; Hep, n=17; Hep+AIP, n=10; * p< 0.05, Holm-Sidak test after one-way ANOVA.). Data are presented as means ± SEMs.

Compared with the control group, fewer APs were evoked in response to depolarizing currents after injection of heparinase (Figure 3A). Analysis of input-output curves showing the averaged number of APs per the intensity of stimulations revealed a significant reduction in cell excitability in the heparinase-treated neurons and restoration of excitability by AIP (Figure 3C). Another indicator of excitability is the spike threshold (Scott et al. 2014). After the heparinase treatment, neurons started to fire at more positive membrane potential in the heparinase group as compared to control and this effect was abrogated by AIP (Figure 3D). Analysis of two peaks in the second derivative of APs, which correspond to AP generation at the AIS and soma [24], revealed a tendency for reduction in the magnitude of the first peak after heparinase treatment (but not the second peak or interval between peaks), and the significant increase of the first peak after CaMKII inhibition, suggesting the modulation of AIS excitability (Figure 3E). An axonal site of heparinase action is also indirectly suggested by the absence of heparinase effects on the peak spike voltage (AP amplitude) that represents an indicator of somatic sodium channel availability (Figure 3F). Heparinase also reduced, in a CaMKII-dependent manner, the half-width and decay of the action potentials, suggesting some modulation of potassium channels (Figure 3G).

### 3.4. Increased activity of CaMKII and impaired expression of AnkG at the axon initial segment after heparinase treatment are abrogated by AIP

Our previous study revealed that the removal of HSHSs reduces AnkG expression at the AIS *in vitro* and *in vivo* [19]. As we in the present study observed the increased autophosphorylation of CaMKII after heparinase treatment, we investigated if the reduction in AnkG at AIS correlates with changes in CaMKII phosphorylation at the same subcellular domain and whether the pharmacological inhibition of the CaMKII autophosphorylation with AIP could abrogate the effects of heparinase treatment on AnkG expression. To facilitate the quantitative analysis of protein expression in the AIS, it was done *in vitro* as previously described [19]. We observed an increased level of pCaMKII at the AIS after digesting HSHSs, which was reduced by AIP to the control levels (Figure 4). Similar to our previous findings, the removal of HSHSs reduced the expression of AnkG along the 40 μm distance of the AIS relative to the control. In line with our electrophysiological recordings, co-incubating hippocampal neurons with heparinase and AIP restored the expression of AnkG to the control levels (Figure 4). Together with electrophysiological data, these results suggest that the reduction of AnkG expression at the AIS and reduced neuronal excitability after cleaving HSHSs are induced by the increased autophosphorylation of CaMKII.

**Figure 4.**
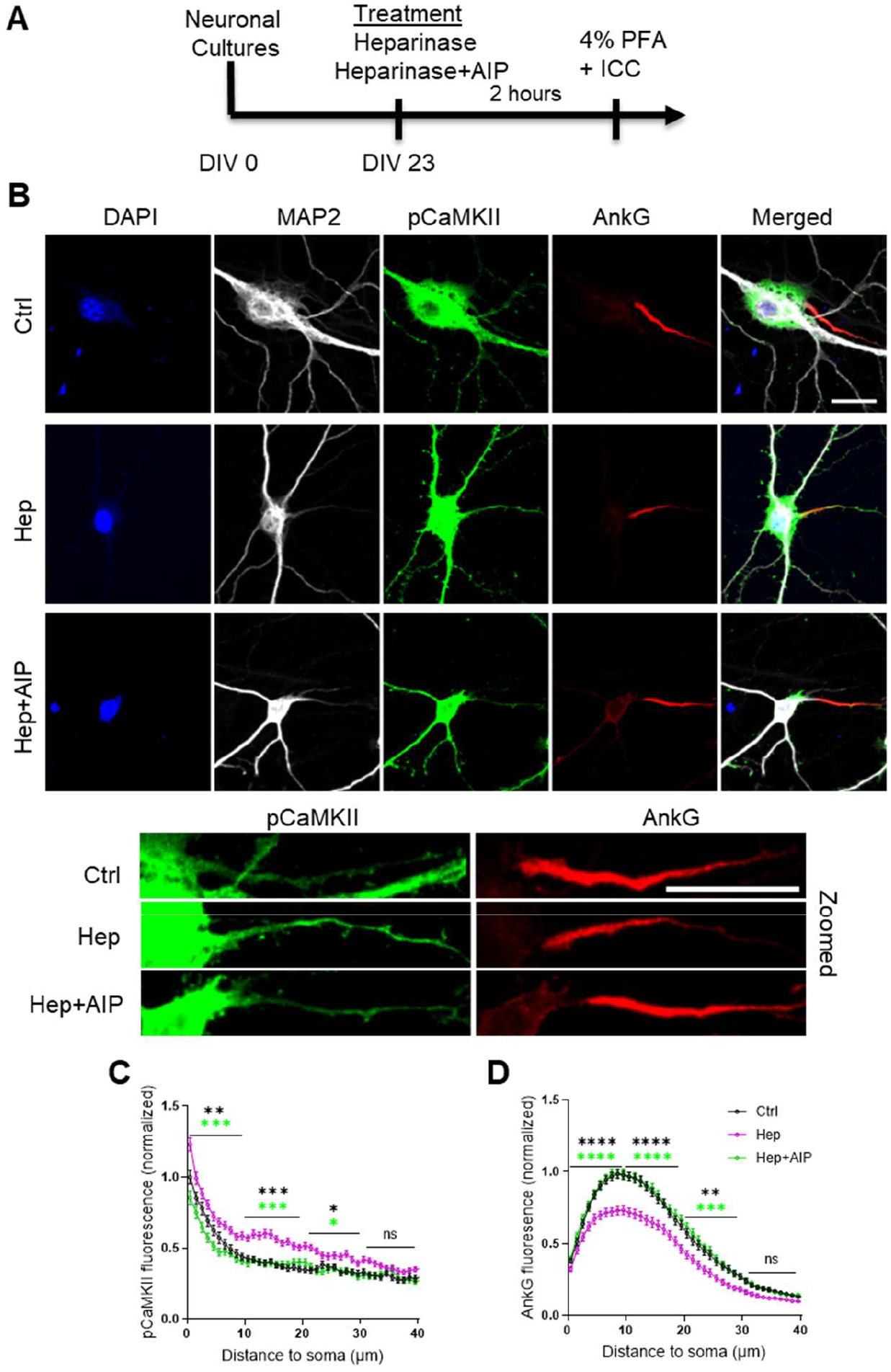
The CaMKII inhibitor AIP abrogates heparinase-induced reduction in AnkG expression in the AIS of hippocampal neurons. (**A**) Timeline showing the treatment of hippocampal neurons with heparinase and AIP for 2 hours followed by immunocytochemistry (ICC). (**B**) Hippocampal neurons were stained with DAPI (blue), MAP2 (gray), pCaMKII (green), and Ankyrin G (AnkG, red) antibodies. Scale bars for both original and zoomed images are 20 μm. (**C & D**) The pCaMKII (C) and AnkG (D) distributions at the AIS were determined by computing a 40 μm-long fluorescence intensity profile with a thickness of 3 pixels in the middle of AnkG-immunopositive areas starting from the edge of the soma (determined using the MAP2 signal). The average expression of AnkG and pCaMKII along the 40 μm line profile of the AIS were quantified from 165 (Ctrl), 183 (hep), and 167 (Hep+AIP) neurons from three independent experiments. Line profiles were normalized to the peak AnkG and pCaMKII values from the control group for each independent experiment. *p<0.05, **p<0.01, ***p<0.001, ****p<0.0001, Holm-Sidak posthoc test after two-way RM ANOVA for the line profiles binned with 10 μm interval.

### 3.5. Impaired recall of contextual memories after heparinase treatment is rescued by coadministration of AIP

In our previous study, we found that heparinase injected before contextual fear conditioning did not affect the level of spontaneous freezing/immobility before conditioning but impaired context discrimination 24 hours after conditioning [19]. This experiment, however, did not allow for discrimination on whether HSHSs are essential for the acquisition, consolidation or recall of contextual memories because reexpression of glycans is a slow process taking several weeks [25] and hence the removal of HSHSs before conditioning would result in impaired HSHS expression during acquisition, consolidation and recall of memories next days after conditioning. In the present study, we specifically tested if HSs are necessary for proper contextual memory recall by injecting heparinase on day 2 after CFC (Figure 5a), i.e. after the acquisition and consolidation of memories were successfully completed. This was confirmed by normal freezing time in the conditioned context and normal context discrimination on day 1 in mice pre-assigned to all experimental groups, i.e. control, heparinase, and heparinase plus AIP (Figure 5a). Also, on day 3 after CFC, i.e. 24 hours after heparinase injection, the freezing time in the conditioned context was normal in the control group, but heparinase-treated mice showed increased freezing in the neutral context (Figure 5B) and impaired contextual discrimination (Figure 5C). Coadministration of AIP restored normal context discrimination after heparinase treatment, not affecting freezing time in the conditioned context.

**Figure 5.**
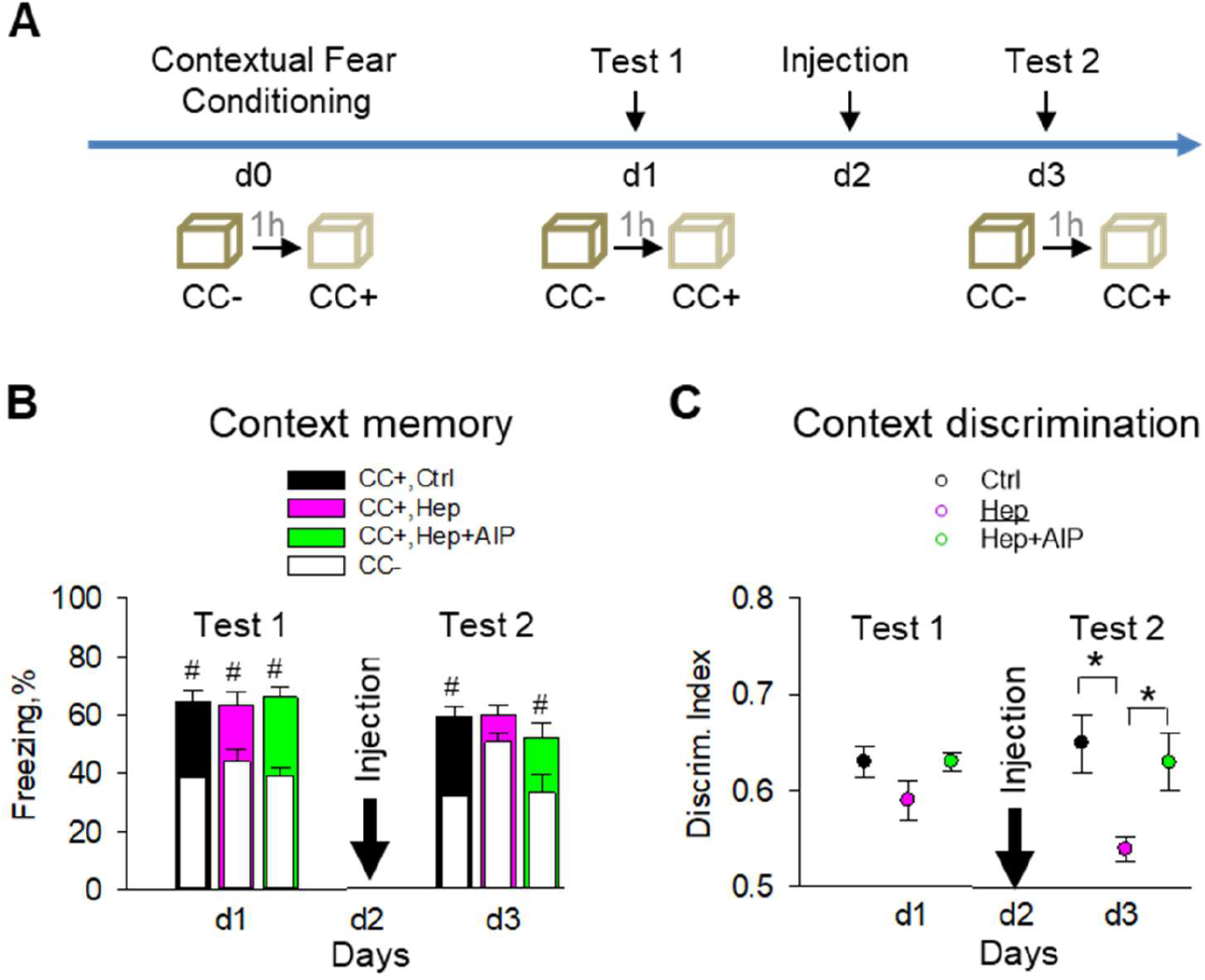
CaMKII inhibitor AIP rescues impaired context discrimination after heparinase injection *in vivo*. (**A**) Scheme of the experiment. (**B & C**) Intrahippocampal heparinase injection 24 h after fear conditioning increases freezing in the neutral context (B) and impairs context discrimination (C), which can be rescued by inhibition of CaMKII with AIP. #P<0.05, paired t-test to compare freezing in conditioned and neutral contexts. *p<0.05, Holm-Sidak posthoc test after two-way RM ANOVA to compare treatments.

## 4. Discussion

Our data show that enzymatic removal of HSHSs in the CA1 region of the hippocampus does not affect miniature postsynaptic currents but leads to reduced axonal excitability of pyramidal cells and impaired contextual discrimination, which correlate with increased activity of CaMKII in general, but particularly in the AIS. Inhibition of CaMKII with AIP normalizes excitability and expression of AnkG in the AIS after heparinase treatment, suggesting a causal link between HSPGs and regulation of axonal excitability via CaMKII autophosphorylation. Below, we discuss the functional importance and possible molecular mechanisms underlying these findings.

Highly expressed in excitatory synapses in the hippocampus, CaMKII has been studied in many aspects of synaptic function such as synaptic strength and synaptic plasticity. Overexpression of α and β isoforms of CaMKII in cultured neurons has opposing effects on mEPSCs strength frequency while CaMKIIβ overexpressing cells exhibit an increase [26]. Thus, it is plausible to assume that in our experiments the effects of increased CaMKIIβ activation were counterbalanced by increased activity of synaptic CaMKIIα, but also we cannot exclude the saturation of CaMKIIβ effects under *in vivo* conditions of the present experiments.

The autophosphorylation of CaMKII, especially of CaMKIIα, on the other hand, has been shown to reduce the excitability of CA1 neurons, which may impact learning [27,28] while inhibition of autophosphorylation of CaMKIIα by a point mutation at T286A increased CA1 neuron excitability. These data are in line with our finding that autophosphorylation of CaMKII was increased after heparinase treatment in the AIS, while expression of AnkG was impaired but could be rescued by the AIP co-administration. Studies show that AIP specifically inhibits CaMKII relative to other kinases such as protein kinase C (PKC), CAMKI and CaMKIV in rat brain extracts [29,30] and in mice [22,23]. The degree of specificity of AIP effects on CaMKIIα versus CaMKIIβ, however, has not been properly resolved.

The AIS, placed in between axonal and somatodendritic domains, is a key structure for the initiation of action potential firing. AnkG, NrCAM, βIV-spectrin, and voltagegated sodium and potassium channels are major structural/functional components of the AIS and their alteration affects AIS assembly and function [31]. βIV-spectrin, an AnkG interaction partner, serves as a bridge between AnkG and actin-based cytoskeleton. Accordingly, animal models harboring AnkG gene deficiency exhibit abnormal animal behavior (such as ataxia) and neuronal excitability in the cerebellum, due to the mislocalization of sodium channels [32]. Progressive ataxia and tremors are also observed in different βIV-spectrin mutant mice (qv^3J^ and βIV null mice) [33]. Findings of the mislocalization of sodium channels in AIS of cerebellar and hippocampal neurons in these mutant mice suggest that altered sodium channel expression is responsible for neurological phenotypes of the mutants. In the cardiomyocyte, βIV-spectrin interaction with CaMKII leads to sodium channel phosphorylation via βIV-dependent targeting CaMKII [34]. Abnormal kinetics of sodium channels and altered cellular excitability after a loss-of-function mutation in the βIV-spectrin gene in the qv^3J^ mouse line suggest that βIV-spectrin/CaMKII complex is an important component for Na^+^ channel regulation in cardiomyocytes. Interestingly, CaMKII is colocalized with βIV spectrin also in AIS of cerebellar Purkinje neurons and qv^3J^ mutant mice exhibit a relatively weak immunostaining signal of CaMKII in AIS of Purkinje cells, implying that βIV-spectrin/CaMKII complex would strongly affect cellular excitability in both heart and brain [34]. Thus, further studies are warranted to study the distribution of βIV-spectrin and ion channels in AIS after targeting of HSHSs.

Extracellularly, the secreted protein gliomedin is a key component at the nodes of Ranvier in the peripheral nerves. The deposition of gliomedin multimers at the nodal gaps facilitates the clustering of the axonodal cell adhesion molecules neurofascin and NrCAM and sodium channels by binding to HSPGs [35]. In cortical neurons, agrin binds to a tyrosine kinase receptor which results in the elevation of intracellular Ca^2+^ and subsequent activation of CaMKII signalling pathway [36]. Regarding potential protein carriers of HSHSs responsible for the regulation of CaMKII activity at the AIS, there are several candidates: Glypicans 1 and 2 are expressed axonally [37,38]. Glypican-4 is also enriched on hippocampal granule cell axons and can bind to its partner orphan receptor GPR158 [39]. Additionally, syndecans are known to be localized at the nodes of Ranvier [40] and axons [3,41]. Syndecans 2 and 3 can directly bind to CASK/LIN-2 protein via the PDZ domain [4] that regulates CaMKII activity in neurons [42]. Further studies of AIS in mice deficient in these HSPGs could be instrumental to identify their role in AIS assembly and axonal excitability via regulation of CaMKII.

Our behavioral experiments for the first time suggest the role of HSHSs in the proper recall of contextual memories and show that *in vivo* inhibition of CaMKII by AIP could abrogate the hypergeneralization induced by heparinase. Previously, AIP has been shown to significantly protect neurons from NMDA-induced neurotoxicity [43], fully restore contractility in cardiac muscles of diabetic rats [44], inhibit doxorubicin-induced apoptosis of cardiac cells [45] and prevent the reinstatement of morphine-seeking behavior in rats [46]. As hypergeneralization is common for several conditions [47-49], targeting this mechanism might be of therapeutic value.

A similar loss of context discrimination as found after heparinase treatment occurs when contextual memories are transferred from the hippocampus to the anterior cingulate cortex via the retrosplenial cortex. Moreover, high-frequency stimulation of memory engrams in the retrosplenial cortex one day after learning produced a recent memory with features normally observed in consolidated remote memories, including contextual generalization and decreased hippocampal dependence [50]. Thus, our data can be interpreted as that a recent contextual memory is distributed in several brain areas and if the hippocampal engrams, in particular CA1, are not activated enough due to a loss of excitability induced by heparinase, another, presumably cortical, representation is used.

In summary, our data make a stronger link between HSHSs and regulation of neuronal excitability and implicate CaMKII in this regulation. Aberrant expression or activity of HSPGs is associated with some pathological conditions like glioblastoma, Fragile X syndrome, neuroinflammation and Parkinson ‘s disease [51-54]. Also, HSPGs are known to bind and co-aggregate with Aβ [55,56]. In light of reported neuronal hyperexcitability in Alzheimer ‘s patients and models of Alzheimer ‘s disease [57,58], our work suggests that Aβ-HSPG interactions may affect the expression of HSPGs at the AIS, decreasing activation of CaMKII at AIS and hence increasing neuronal excitability. At synapses, Aβ is known to inhibit autophosphorylation of CaMKII at Thr286 and impair synaptic plasticity [59]. Thus, our study suggests potential pathophysiological mechanisms and indicates an option to prevent these by targeting CaMKII signaling at AIS.

## Author Contributions

Conceptualization, Hussam Hayani and Alexander Dityatev; Data curation, Michael Druzin; Formal analysis, Inseon Song, Tatiana Kuznetsova, David Baido-Ansah, Hadi Mirzapourdelavar, Oleg Senkov, Hussam Hayani and Andrey Mironov; Funding acquisition, Staffan Johansson and Alexander Dityatev; Investigation, Inseon Song, Tatiana Kuznetsova, David Baido-Ansah, Oleg Senkov, Hussam Hayani and Andrey Mironov; Methodology, Inseon Song, Oleg Senkov, Rahul Kaushik and Michael Druzin; Resources, Staffan Johansson and Alexander Dityatev; Supervision, Inseon Song, Rahul Kaushik, Michael Druzin, Staffan Johansson and Alexander Dityatev; Visualization, Tatiana Kuznetsova and David Baido-Ansah; Writing – original draft, Inseon Song, David Baido-Ansah and Alexander Dityatev; Writing – review & editing, all co-authors.

## Institutional Review Board Statement

All treatments and behavioral procedures were conducted in accordance with ethical animal research standards defined by German law and approved by the Ethical Committee on Animal Health and Care of the State of Saxony-Anhalt, Germany, the license numbers 42502-2-1159 and -1322 DZNE.

## Informed Consent Statement

Not applicable

## Data Availability Statement

Data supporting reported results can be obtained upon a request to the corresponding author (A.D.).

## Acknowledgments

We thank Katrin Boehm for her technical support.

